# A Novel Droplet Digital PCR Human mtDNA Assay for Global Fecal Source Tracking

**DOI:** 10.1101/794891

**Authors:** Kevin Zhu, Brittany Suttner, Konstantinos T. Konstantinidis, Joe Brown

## Abstract

Human mitochondrial DNA (mtDNA) provides a promising target for microbial source tracking because it is unique to humans and universal across human individuals. We developed a droplet digital PCR (ddPCR) assay, hCYTB484, targeting the cytochrome *b* gene of the human mtDNA and compared the performance of the hCYTB484 assay with a widely used assay targeting human-associated *Bacteroides*, the HF183/BacR287 assay. We also defined and validated the analytical limit of detection and analytical lower limit of quantification; these analytical limits determine the concentration levels above which samples are declared to be positive and quantifiable for the target, respectively. We found both assays to be highly specific (95%) against cow and pig feces; however, the hCYTB484 was more sensitive when tested against individual human feces from US, Bangladesh, and Mozambique (100% versus a mean of 51% across the 3 countries). To further compare the performance of the two assays, we utilized a receiver operating characteristic curve, showing that the hCYTB484 marker was widely distributed across human feces from the 3 different geographical regions tested and in higher quantities than the HF183/BacR287 marker. The higher performance of the hCYTB484 marker in individual feces is a desirable characteristic in the detection of fecal pollution from sources to which fewer individuals contribute, such as non-sewered types of sanitation that serve most of Earth’s population and carry the highest risk of exposure to fecal-oral pathogens.

**Importance:** The usefulness of an ideal human-specific, culture- and library-independent marker to the microbial source tracking field is reflected by the numerous efforts to develop such markers; however, thus far, microbial-based markers of this type have exhibited variable source-specificity across geographies and variable sensitivity across scales of fecal waste. Most of the world’s population is served by non-sewered forms of sanitation that also carry the highest risk of exposure to fecal-oral pathogens. This reality underscores the need for markers of human fecal contamination that have high sensitivity in fecal pollution sources to which fewer individuals contribute to, such as fecal sludges found in pit latrines. We show that human mtDNA-based methods can be highly source-specific and highly sensitive in smaller scales of fecal pollution, providing a useful addition to the microbial source tracking toolbox by complementing the variable performance of microbial-based markers.

## Introduction

Within the microbial source tracking (MST) field, the development of an ideal human-specific, culture- and library-independent marker has been a goal for many research efforts for a variety of reasons—notably, its potential as a relatively rapid and broadly-applicable method not requiring the extent of local validation of the host and marker relationship that is required in library-dependent methods (1–4). However, MST marker studies have demonstrated variable specificity (true negative detection rate, calculated by the number of true negatives divided by the sum of true negatives and false positives) and sensitivity (true positive detection rate, calculated by the number of true positives divided by the sum of true positives and false negatives) performances for human-associated microbial DNA markers across varying geographies (1, 5–8), possibly influenced by variations in the human gut microbiome (9) and human-animal interactions (5, 6). Microbial DNA markers have also exhibited variable performance across different scales of fecal pollution: high sensitivity in wastewater collected from sewage networks but lower sensitivity in human fecal samples (10). While much effort has been made in the development of human-specific, culture- and library-independent markers, the variable performance of such microbial markers across various settings necessitates their local validation in the intended setting prior to use to ensure that the fecal signal can be unambiguously attributed to a source (6, 8, 11).

Most of the global human population does not have access to sewered sanitation: an estimated 56% of households globally have unsewered sanitation facilities and 12% of households globally have no sanitation facility (12). Furthermore, disproportionalities in sanitation exist: the same geographic regions that contain the lowest rates of sewered households are where most of the human population live and where the most human feces are produced (12). This means that for much of the world’s population, enteric pathogen exposure risks are related to sludges and other non-wastewater fecal waste streams that are composites of feces from relatively few individuals. This reality underscores the need for markers to serve in situations where exposure to fecal sludge is more relevant than exposure to wastewater collected through a sewerage network. In these settings, markers that are widely distributed throughout humans are desired because with fewer individuals contributing to fecal sludges, the presence of the marker in every individual is crucial.

Mitochondrial DNA (mtDNA) based methods differ principally from microbial-based MST methods in that they target host DNA directly rather than host-associated microbial DNA. Because mtDNA methods target host DNA conserved across the species, the link between host and fecal indicator microbe is no longer relevant, and, therefore, mtDNA methods do not require the same extent of validation as emphasized for host-associated, microbial DNA markers. Additionally, because mtDNA is universally distributed across human individuals, mtDNA methods may offer superior sensitivity in individual feces and smaller scales of fecal pollution. Quantitative human mtDNA-based assays for MST have exhibited high specificity in the US (13, 14) and in China (15); however, mtDNA-based methods are found in lower numbers in sewage samples relative to microbial-based MST markers (14, 15).

Because mtDNA-based markers are typically found at lower concentrations than microbial-based makers, we utilized the droplet digital PCR (ddPCR) platform for its improved reproducibility in lower concentrations. As a tool for MST, ddPCR has demonstrated several appealing characteristics when compared to qPCR, including greater tolerance of PCR inhibitors (16), reduced quantification bias when duplexing assays (16) and improved reproducibility over qPCR (17, 18). While implementing previously-designed human mtDNA assays (13, 14) on the ddPCR platform, we observed notable noise when differentiating between positive and negative partitions. One assay was implemented as a mismatch amplification mutation assay by intentionally designing penultimate primer mismatches to increase species-specificity (13), potentially inducing PCR inefficiencies and leading to increased noise on ddPCR (19). Another assay (14) contained a probe that spanned several population-variable polymorphisms (20–24). Because the mitochondrial genome is maternally inherited, polymorphisms have been studied for their utility in identifying human individuals from different matrilineages. Several studies have investigated the use of polymorphisms within the cytochrome b gene to discriminate between populations of people, finding that combinations of polymorphisms can be used to discriminate between different populations of people (20, 21). Because mismatches between template and oligonucleotides can result in inefficiencies in the PCR (19, 25), the discriminatory power of these polymorphisms alludes to the potential for geographic variations in performance of mtDNA-based methods due to population-specific polymorphisms. Assay PCR inefficiencies in ddPCR potentially increase the noise surrounding the raw signal, resulting in potentially variable behavior on the ddPCR platform.

In this study, we developed a novel assay (hCYTB484) targeting human mtDNA on the ddPCR platform with three goals: 1) specificity to human mtDNA, 2) optimization for ddPCR conditions, and 3) avoidance of identified polymorphisms for a geographical stable assay. We compared the performance of our mtDNA assay with that of a high performing assay targeting human-associated *Bacteroides*, the HF183/BacR287 assay, in parallel on ddPCR. The HF183/BacR287 assay (2) is an improved version of a widely-studied microbial DNA assay (26). To investigate the geographic variability of the two assays in situations where few individuals contribute to the fecal pollution, we tested the assays on human feces obtained from the US, Bangladesh, and Mozambique. We defined and tested the performance characteristics of each assay, including sensitivity, specificity, analytical limit of detection (aLoD, the limit above which samples are considered positive for the target) and analytical lower limit of quantification (aLLoQ, the limit above which samples are considered quantifiable for the target). We then compared sensitivity and specificity performances between the two assays by using receiver operating characteristic (ROC) curves to detail performance beyond scalar values of sensitivity and specificity. We hypothesized that by targeting a widely conserved host-mtDNA marker, we would develop an assay that is more sensitive in human feces from various geographies than the microbial-based HF183/BacR287 assay.

## Results

### Analytical limit of detection

The ddPCR platform discretizes a 20 microliter PCR reaction into nanoliter-scale reactions, referred to here as partitions, by emulsifying the PCR reaction within oil droplets. Each ddPCR well contains tens of thousands of such partitions. After thermal cycling the wells, the ddPCR platform reads a presence/absence signal from each partition in a ddPCR well and calculates a concentration from the ratio of the number of partitions that were read as positive (positive partitions) to the total number of partitions from a ddPCR well that were read (accepted partitions). The formula (QuantaSoft Version 1.7.4.0917) used to estimate the concentration of target in the original sample can be written as:

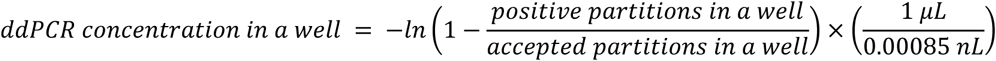

From the above equation, the number of positive partitions and accepted partitions represent the raw signal of the ddPCR platform. The ddPCR concentration in a well represents the number of copies of the target found in the 20 microliter PCR reaction; we will refer to copies of target per ddPCR well, which is the ddPCR concentration in a well described above multiplied by 20 μL.

To assess false-positive rates for both the hCYB484 and HF183/BacR287 assays on ddPCR, we analyzed UV-treated, molecular-grade water. While approximately 95% of analyzed blanks lacked any amplification (as defined by any number of positive partitions greater than zero) for both assays, our results showed that false-positive partitions occurred at low concentrations and low rates for both assays (Table 1). We chose to define our analytical limit of detection (aLoD) as the minimum level of the analyte in a sample that will be reported as detected with 95% frequency across replicates of ddPCR wells. Prior to performing experiments to validate the aLoD, we used the Poisson distribution to calculate an expected value of 3 positive partitions per ddPCR well as the level of concentration within the ddPCR well that would enable us to observe at least 1 positive partition in 95% of ddPCR wells. In our experiments to validate an aLoD of 3 positive partitions per well, we achieved mean positive partitions per well of 4.54 (hCYTB484) and 4.83 (HF183/BacR287). We compared sampling distributions from our ddPCR aLoD experiments with corresponding expected Poisson distributions using the Anderson-Darling test and obtained *p*-values of 0.85 and 0.98 for the hCYTB484 and HF183/BacR287 assays, respectively (Figure S1).

**Table 1.** Results of analyzing no-template controls (UV-treated molecular-grade water) on ddPCR for both assays.

| Number of Positive Partitions in a Well | Frequency of Occurrence |  | Relative to aLoD |
| --- | --- | --- | --- |
|  | HF183/BacR287<br>(n = 94) | hCYTB484<br>(n = 93) |  |
| 0 | 97.85 % | 95.74 % | Below aLoD |
| 1 | 2.15 % | 3.19 % |  |
| 2 | 0.00 % | 0.00 % |  |
| 3 | 0.00 % | 0.00 % | Above aLoD |
| 4 | 0.00 % | 1.06 % |  |

### Analytical lower limit of quantification

To determine the analytical lower limit of quantification (aLLoQ), we defined our aLLoQ goal as the level of concentration that would result in a coefficient of variation (CV, calculated by the ratio of the standard deviation to the mean) of the copies per ddPCR well equal to 25%. The coefficient of variation and the relative standard deviation (RSD) both refer to the same measure. We assayed a variety of dilutions of positive controls at different levels of concentrations and calculated CV values using a relatively large number of ddPCR well replicates (46 technical replicates) (Figure 3). We interpolated the level of concentration that would allow us to achieve a CV of 25% by fitting a linear regression to log_10_ transformed CV and copies per ddPCR well values. The regression fitted for each assay used in this study incorporates some intraassay variability as the 4 estimates of the CV for each assay are from 3 separate serial dilutions created on different days. The linear regressions produced aLLoQs of 28.2 and 29.6 copies per ddPCR well for hCYTB484 and HF183/BacR287, respectively.

**Figure 1.**
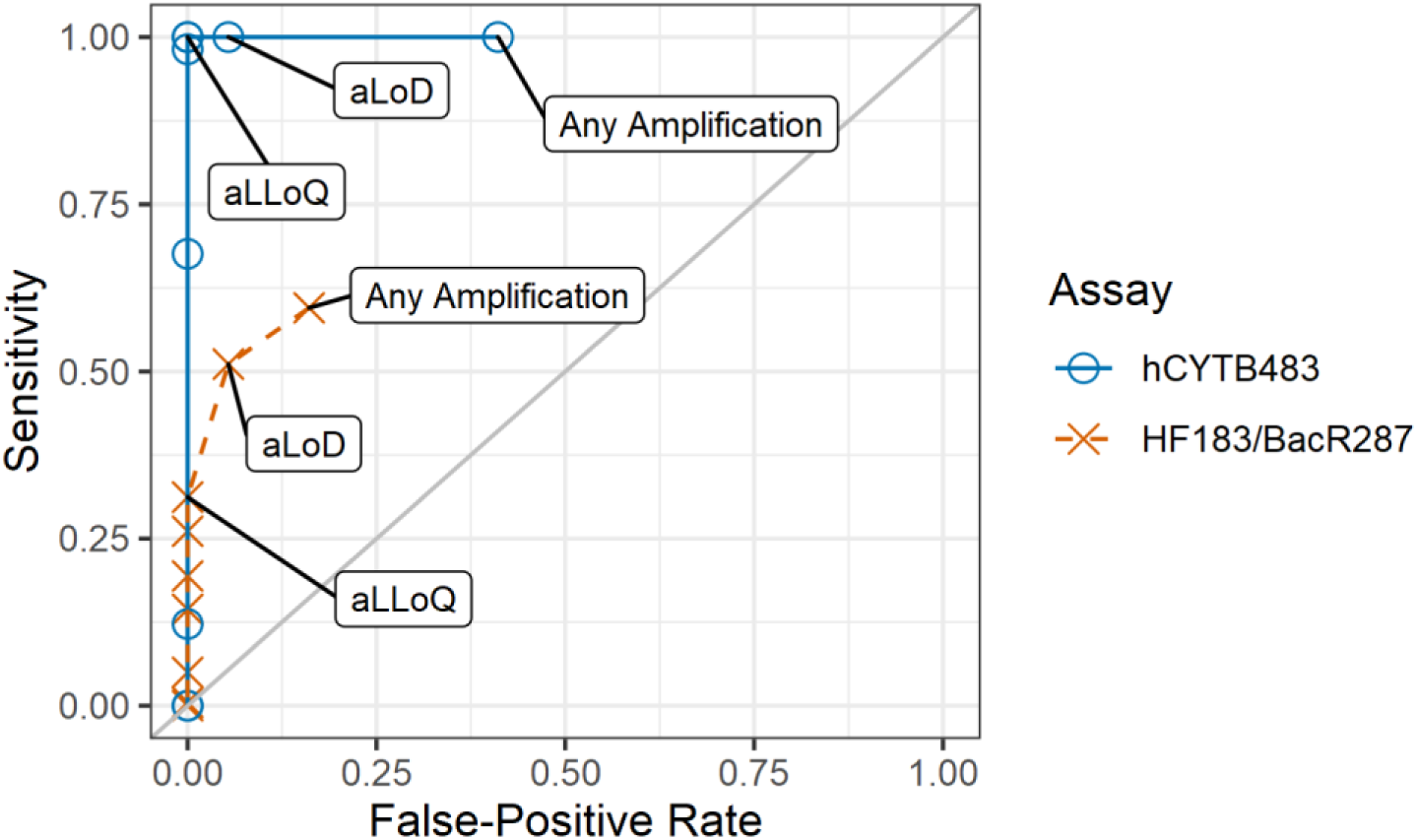
Receiver operating curves for hCYTB484 and HF183/BacR287 ddPCR assays using human (n = 222) and non-human feces samples (n = 56) showing rates of sensitivity and false-positives calculated for a range of thresholds beginning with any amplification, analytical limit of detection (aLoD), and analytical lower limit of quantification (aLLoQ).

**Figure 2.**
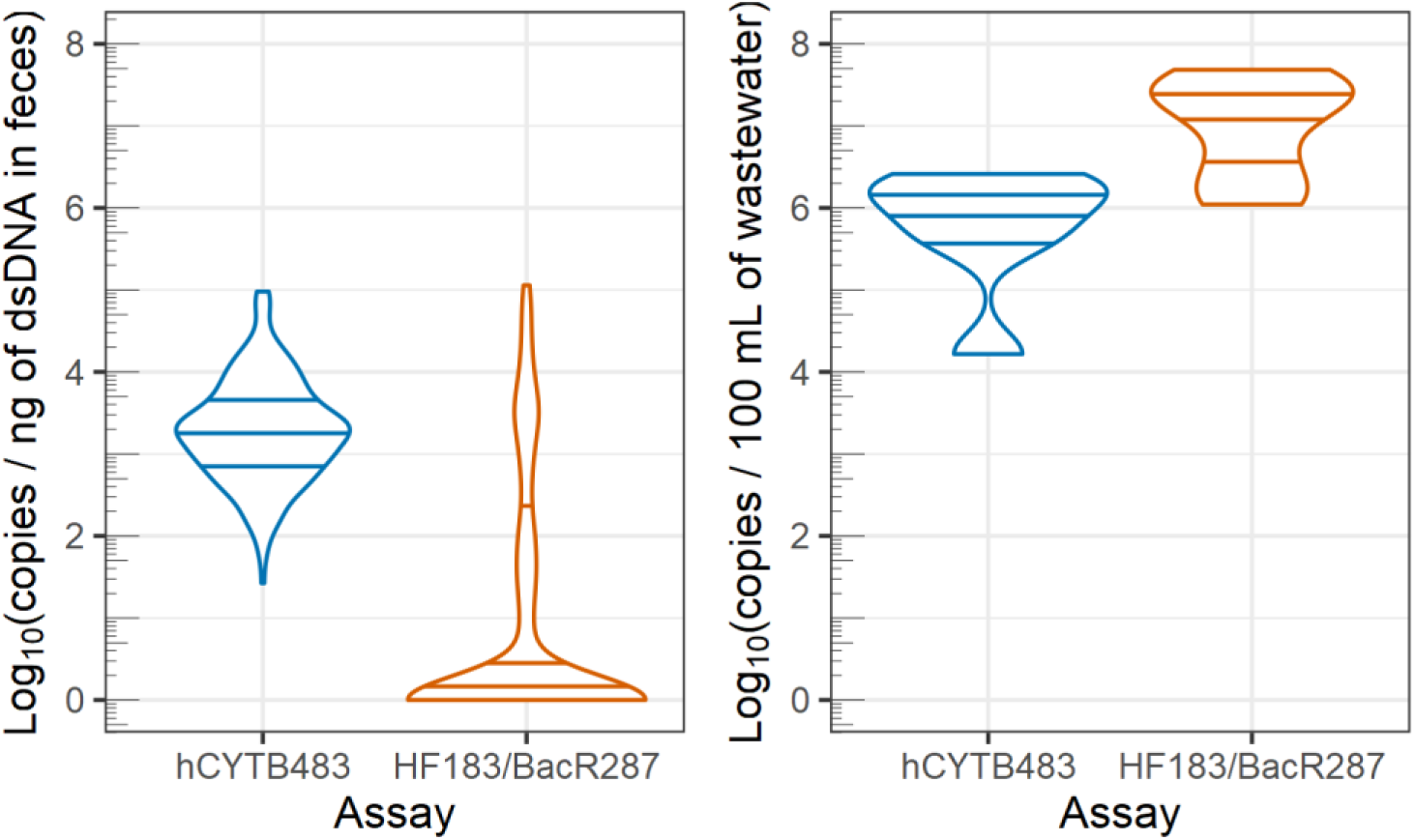
Violin plots of marker concentrations in human feces (left) and human wastewater (right). Concentrations are plotted as log*10*(concentration + 1), and concentrations from feces are normalized to concentration of dsDNA to account for variations in moisture content (ng of dsDNA determined by Qubit). Horizontal lines indicate 25^th^, 50^th^, and 75^th^ quantiles.

**Figure 3.**
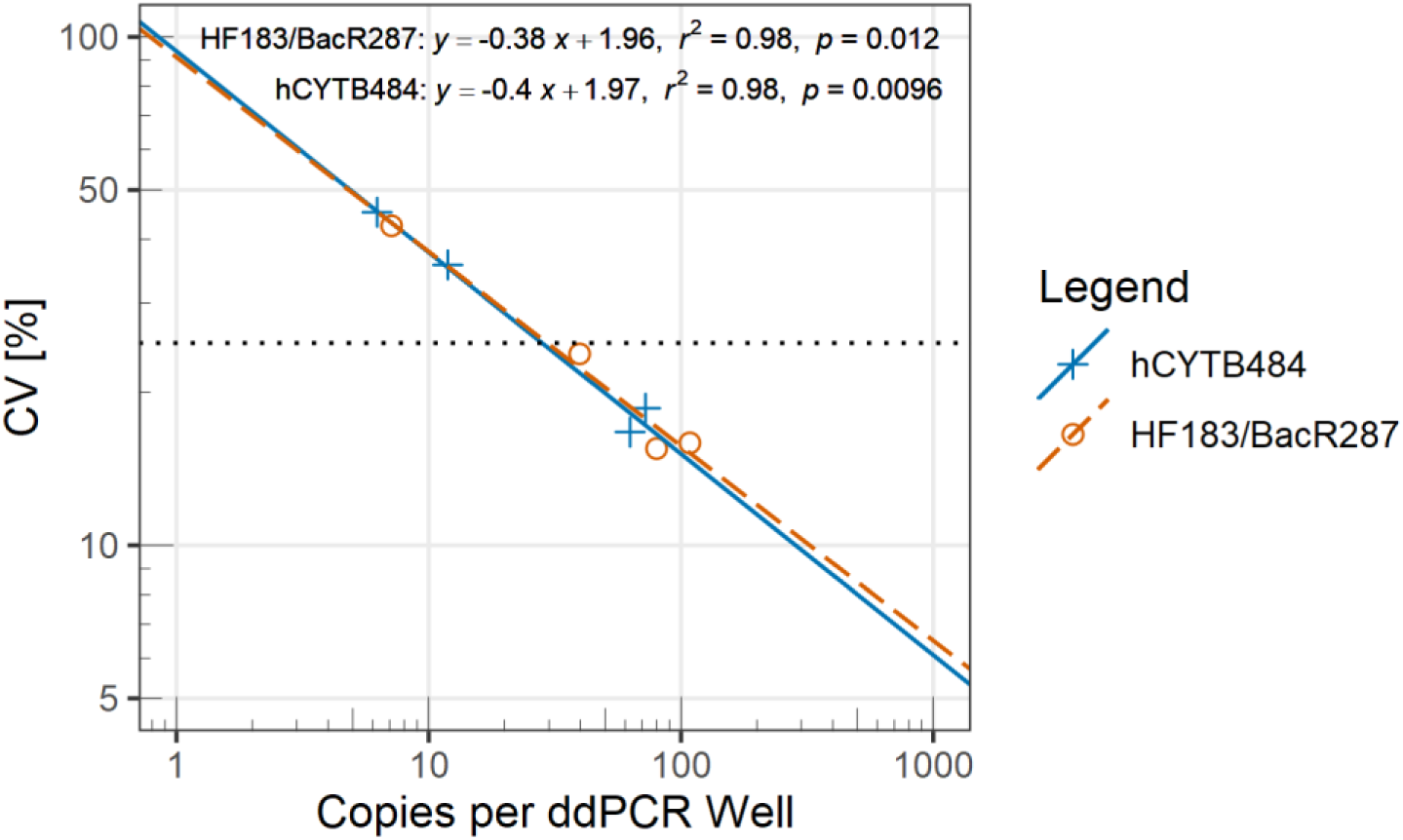
CV values plotted against copies of per ddPCR well for hCYTB484 (simplex) and HF183/BacR287 (multiplex with accompanying IAC) on log_10_-log_10_ axes. We obtained CV estimates for each assay from 4 dilutions made from 3 separate serial dilutions (for each assay), each created on 3 separate days. We fitted linear regressions to the log_10_ transformed values of both CV and copies per ddPCR well (linear equations, *r*^2^ values, and F statistic *p*-values displayed in top right corner of plot). The horizontal dotted line depicts our aLLoQ imprecision target, CV = 25%.

### Sensitivity and specificity

We tested the sensitivity and specificity of HF183/BacR287 and hCYTB484 on feces samples from 56 unique farm animals (cows and pigs) and 222 unique humans (Table 2). We calculated sensitivity and specificity values for these samples over a range of thresholds (Table 3). As the threshold, or the level of concentration at which a well is considered to be positive, is increased from any amplification to the aLLoQ, the hCYTB484 assay exhibited an increase in specificity from 59% to 100% while sensitivity remained constant at 100%. Across the same threshold levels, the HF183/BacR287 assay exhibited an increase in specificity from 84% to 100% but also a decrease in sensitivity from 59% to 32%. The hCYTB484 assay exhibited 100% sensitivity and 100% specificity with the threshold at the aLLoQ, whereas the HF183/BacR287 assay exhibited 100% sensitivity and 32% sensitivity at the same threshold.

**Table 2.** Background information on the human and non-human fecal samples used in this study and results of specificity and sensitivity (with the threshold at which samples are declared positive or negative set at the aLoD).

|  | Sample source | No. of samples | Sample type | Source information | No. of sample positive<br>(threshold set at aLoD) |  |
| --- | --- | --- | --- | --- | --- | --- |
|  |  |  |  |  | hCYTB484 | HF183/BacR287 |
| Non-human | Cow | 22 | Individual | Farm | 1 (5 %) | 0 (0 %) |
|  | Pig | 34 | Individual | Farm | 2 (6 %) | 3 (9 %) |
|  | Total Specificity = |  |  |  | 95 % | 95 % |
| Human | Feces | 222 |  |  |  |  |
|  | United States | 11 | Individual | Adults | 11 (100 %) | 10 (90 %) |
|  | Bangladesh | 140 | Individual | Children | 140 (100 %) | 60 (43 %) |
|  | Mozambique | 71 | Individual | Children | 71 (100 %) | 44 (62 %) |
|  | Untreated Wastewater | 4 | Composite | 16 MGD to 90 MGD | 4 (100 %) | 4 (100 %) |
|  | Total Sensitivity = |  |  |  | 100 % | 51% |

**Table 3.**
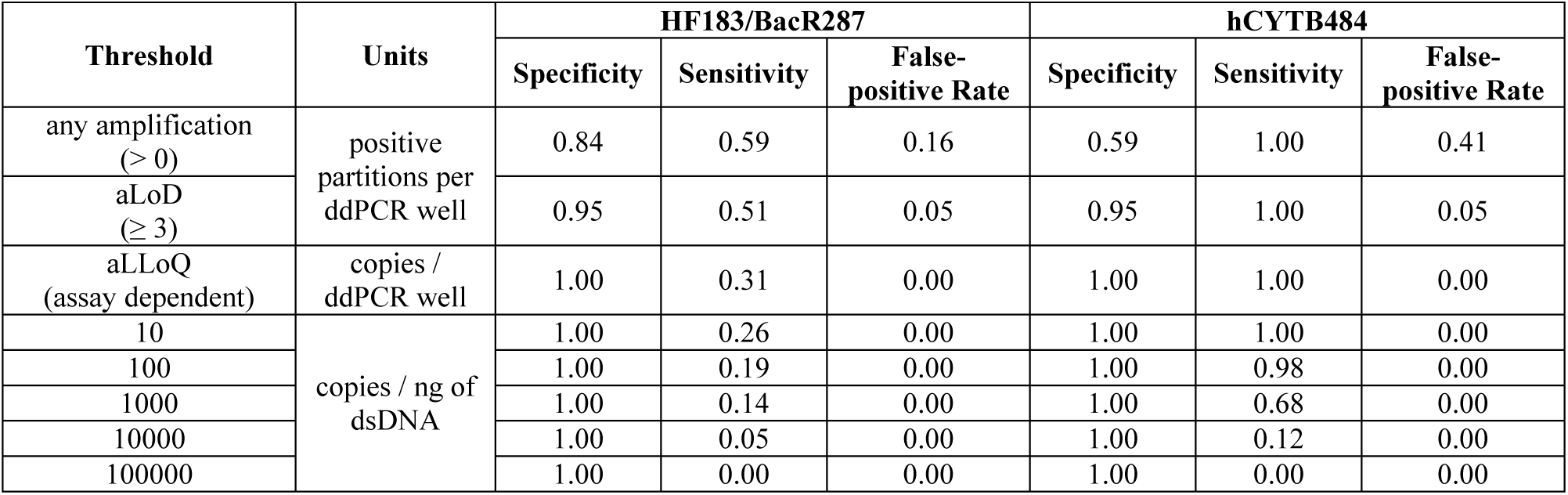
Rates of specificity, sensitivity, and false positives calculated for a range of thresholds for both assays.

| Threshold | Units | HF183/BacR287 |  |  | hCYTB484 |  |  |
| --- | --- | --- | --- | --- | --- | --- | --- |
|  |  | Specificity | Sensitivity | False-positive Rate | Specificity | Sensitivity | False-positive Rate |
| any amplification (> 0) | positive partitions per ddPCR well | 0.84 | 0.59 | 0.16 | 0.59 | 1.00 | 0.41 |
| aLoD ($\geq 3$) | | 0.95 | 0.51 | 0.05 | 0.95 | 1.00 | 0.05 |
| aLLOQ (assay dependent) | copies / ddPCR well | 1.00 | 0.31 | 0.00 | 1.00 | 1.00 | 0.00 |
| 10 | copies / ng of dsDNA | 1.00 | 0.26 | 0.00 | 1.00 | 1.00 | 0.00 |
| 100 |  | 1.00 | 0.19 | 0.00 | 1.00 | 0.98 | 0.00 |
| 1000 |  | 1.00 | 0.14 | 0.00 | 1.00 | 0.68 | 0.00 |
| 10000 |  | 1.00 | 0.05 | 0.00 | 1.00 | 0.12 | 0.00 |
| 100000 |  | 1.00 | 0.00 | 0.00 | 1.00 | 0.00 | 0.00 |

## Discussion

### Analytical limit of detection

Assaying analytical blanks for both assays demonstrated that positive partitions occur even when a no-template control has been analyzed, albeit at low rates (< 5%) and low concentrations (mostly below aLoD). It is possible that the hCYTB484 assay exhibited slightly higher rates of false positive because the assay targets human mtDNA; there are several opportunities for human mtDNA to contaminate the reaction mixture due to the series of manipulations required for ddPCR. Combining the results from the analytical blanks with the false-positive rates from non-human feces, we show that very low concentrations (below the aLoD) of false positives occur at low rates when using the hCYTB484 assay, and, consequently, we recommend that the threshold be set at or above the aLoD and care be taken to avoid contamination of human mtDNA.

The sampling distributions of our aLoD ddPCR experiments were similar to the corresponding theoretical Poisson distributions, which suggests that the Poisson distribution is a representative model for the number of positive partitions per well in the ddPCR process. We utilized the Anderson-Darling test to assess whether each sampling distribution was drawn from a Poisson distribution created using the sampling distribution mean as the maximum likelihood estimator, resulting in *p*-values of 0.85 and 0.98 for the hCYTB484 and HF183/BacR287 assays, respectively (Figure S1). While we used the Poisson distribution as a representative model for the number of positive partitions per well in the ddPCR process, we note that it is a theoretical stochastic model and does not account for potential biases introduced by external sources of variability (27), such as the operator or equipment. Using our findings at the mean positive partitions per well of 4.54 and 4.83, we extrapolated our aLoD down to a mean of 3 positive partitions per well using the Poisson distribution and our target of detection of a theoretical 95% probability. This aLoD of 3 has been used and accepted in both qPCR (28) and ddPCR applications (16).

### Estimation of the coefficient of variation and mean positive partitions per well

Too few replicates can result in an erroneous estimation of the coefficient of variation (CV), as has been noted in qPCR (29). To avoid such erroneous estimations and to better understand how values of the CV change as a function of the level of concentration, we sought to estimate the true values of the CV by using a relatively large number of replicates (*n* = 46). Although the CV/RSD of the target concentration is widely used as an imprecision goal when determining the ddPCR aLLoQ, there is no consensus on the number of replicates required, with 5 to 20 replicates being commonly used (30–32). In the absence of agreement on the number of replicates, we used many replicates in combination with cumulative moving plots to assess our estimation of the mean positive droplets per well and the concentration CV (Figures S2 and S3). By construction, the last point in a cumulative moving plot will coincide with the parameter value calculated from the total number of replicates; however, what is of interest is if the cumulative moving parameter converges before the total number of replicates and remains steady. The cumulative moving mean positive droplets per well and cumulative moving concentration CV converged before the total number of replicates in most cases, indicating that the use of 46 replicates was more than enough. We note that the use of less than 20 replicates did not appear to offer an effective estimation of the true CV as some of the values of the CV based on less than 20 replicates fell outside of the 95% confidence intervals. Compared to 20 replicates, 30 or more allowed for a greater degree of convergence.

### Analytical lower limit of quantification

We chose to set our aLLoQ to the level of concentration that would result in a coefficient of variation (CV, as calculated by the standard deviation of concentrations divided by the mean of concentrations) of 25% (30, 31). While aLLoQs are often set using “less-than or equal to” a CV target (for example, ≤ 25%), the difficulty in finding the level of concentration that results in a specific CV results in potentially setting aLLoQs by satisfying the less than condition (< 25%) rather than the equal to (= 25%) condition. However, this can cause variation in aLLoQs set using the same CV target; hypothetically, aLLoQs that have been set based using levels of concentrations that resulted in a CV of 23% versus a CV of 13% are both valid for an aLLoQ based on ≤ 25%, and the difficulty in repeatedly trying different levels of concentration to achieve a CV of 25% deters attempts to satisfy the “equal to” condition. In an attempt to develop a more standardized approach, we interpolated an aLLoQ using a regression fitted to observed CV and concentration values, referred to as a variance function or imprecision profile.

While variance functions for ddPCR assays have not yet been widely explored, various forms of variance functions have been used in diverse fields (33, 34). Variance functions can have many forms (35). Because the CV is a measure of variance, a scale parameter, we plotted the CVs on a log-scale and plotted concentration on a log-scale and observed a linear relationship between data points (Figure 3). Despite different numbers of replicates used to estimate CV values, we also observed similar linear regressions fitted to the log_10_-transformed CV values and log_10_-transformed copies per ddPCR well (Figure 4) in imprecision profiles published by others (30, 31). In Figure 4, we plotted linear regressions derived from multiple assays (including both simplex and duplex assays) from each referenced study. Investigating differences in imprecision profiles between simplex and multiplex assays was outside the scope of this study; however, the imprecision profile for the HF183/BacR287 assay (multiplexed with an IAC) was similar to the hCYTB484 simplex assay (Figure 3), which is supported by the relative absence of a quantification bias when duplexing on the ddPCR platform (16). Because of the agreement between linear regressions for our data and data published by others, we hypothesize that assays implemented on the Bio-Rad QX100/QX200 ddPCR platforms may share similar imprecision profiles. This line of thought is supported by the fact that ddPCR calculates a concentration from direct counts of many endpoint PCR results, eliminating some variability associated with individual assay performance due to the nature of endpoint PCR. Knowledge of an imprecision profile common to assays implemented on a ddPCR platform can aid operators in standardizing the determination and validation of analytical limits. An interpolation approach to determining an aLLoQ can be helpful in allowing operators to set aLLoQs in a more standardized approach by providing a more precise estimation of the level of concentration that satisfies the aLLoQ definition rather than trying various levels of concentration in attempts to satisfy the less than or equal to target CV. In order to leverage an interpolation approach, we need an appropriate model with which to fit to observed data. The combination of relatively high values of *r*^2^ and low *p*-values (Figure 4) leads us to believe that a linear regression fitted to log_10_ transformed CV values and log_10_ transformed copies per ddPCR well is a potential model for the imprecision profile of assays on the Bio-Rad QX100/QX200 ddPCR platforms. Further research investigating more models, assays, and levels of concentration across more research groups performing ddPCR is warranted.

**Figure 4.**
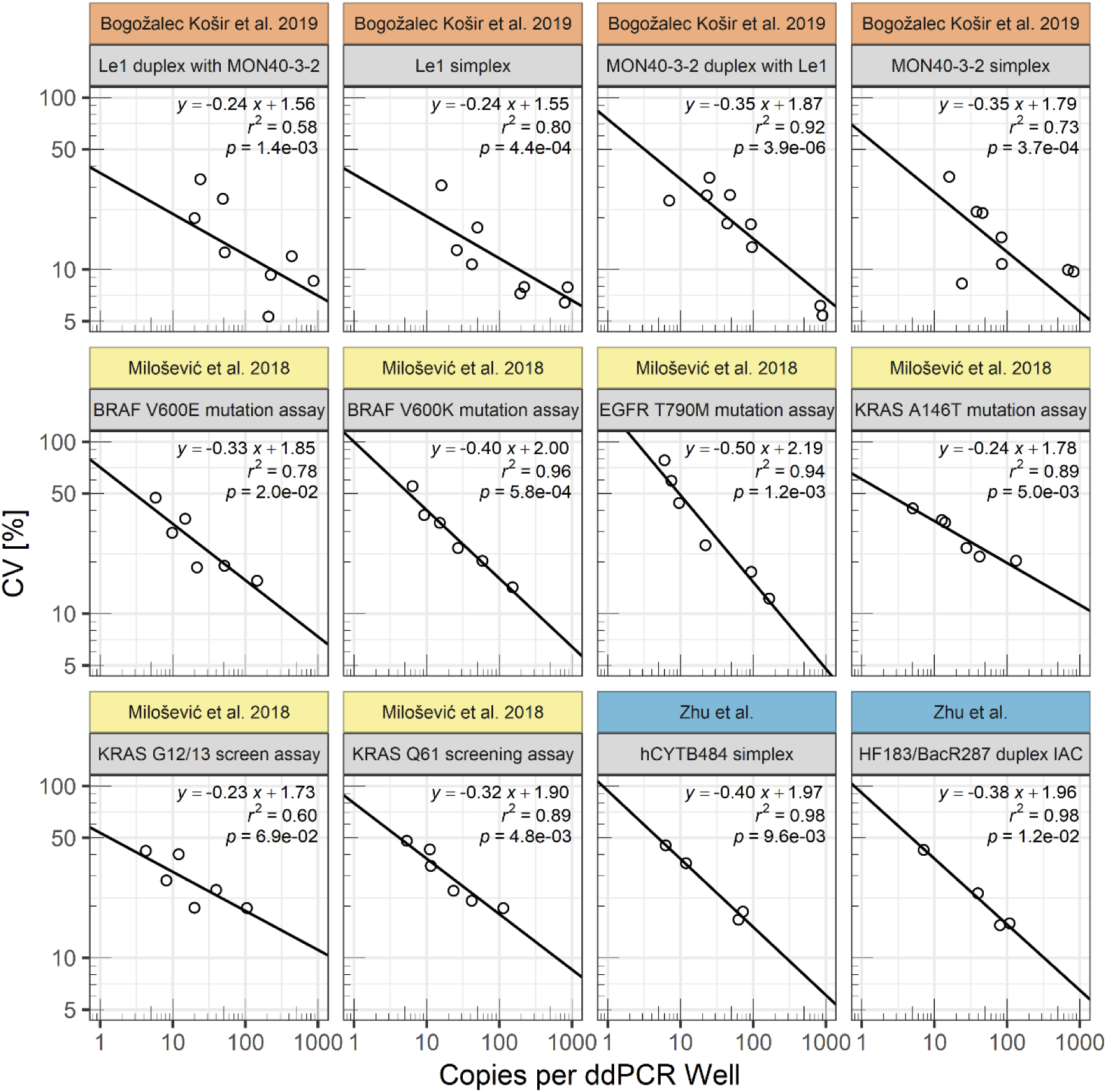
CV values plotted against copies per ddPCR well on log_10_-log_10_ axes for data obtained from this study (Zhu et al.), Bogožalec Košir et al. 2019 (30), and Milošević et al. 2018 (31). We fitted linear regressions to the log_10_ transformed CV values and log_10_ transformed copies per ddPCR well. We report linear equations, *r^2^* values, and F-statistic *p*-values (for the slope coefficient) displayed in top right corner of each plot. The 12 plots include both simplex and duplex assays with assay name indicated under each study label.

### Comparing marker performance

To compare sensitivity and specificity between the two assays, we created receiver operating characteristic (ROC) curves by plotting sensitivity and specificity over a range of thresholds. ROC curves are a useful method of comparing detailed classifier performance because they communicate the dependency of sensitivity and specificity on threshold (36, 37). Communicating the dependency of sensitivity and specificity on threshold is particularly useful because in the case of molecular methods, such as qPCR, different approaches to distinguishing a sample as positive have been used. In some cases, different thresholds for declaring a sample as positive can result in dramatically different sensitivity and specificity performances being calculated with the same data (38). By showing how sensitivity and specificity vary with threshold, ROC curves communicate more detailed performance that may be crucial to selecting a threshold, potentially facilitating standardization of the implementation of assays. For instance, in our ROC curves, we show that the hCYTB484 specificity dramatically decreases from the aLoD to any amplification and has the best specificity at the aLLoQ. This communicates the crucial detail that although the hCYTB484 assay can exhibit low levels of false-positives, these false-positives can be avoided by using the aLoD or aLLoQ as the threshold, with the aLLoQ offering a very high specificity performance. Assays targeting the HF183 human-associated *Bacteroides* sequences have been shown to exhibit some of the weaknesses of the microbial DNA based markers as discussed in the introduction (6, 8, 10, 11). Considering these findings, researchers have proposed that it may be beneficial to utilize HF183 markers in conjunction with other markers to address its weaknesses in the so-called MST “toolbox” approach. ROC curves offer a potentially useful method for assembling such a toolbox by communicating and comparing detailed assay performance to investigators looking for assays to fit their needs.

In comparison with previously published qPCR human mtDNA assays, our hCYTB484 ddPCR assay detected greater concentrations (copies / 100 mL of wastewater) of human mtDNA in both human feces and wastewater (13, 14). However, we still detected the hCYTB484 marker at lower concentrations than the HF183/BacR287 marker in wastewater (Figure 2). Compared to the HF183/BacR287 marker, the hCYTB484 marker maintains higher sensitivities at all levels of threshold in human feces (Table 3). Through the ROC curve, we demonstrated the hCYTB484 marker to be widely distributed in individual human feces in concentrations above the aLLoQ, indicating that the hCYTB484 marker is potentially useful in detecting human fecal sludges and other concentrated fecal waste streams where fewer individuals contribute to the waste. Such matrices are the most common form of fecal wastes globally where risks of exposure are highest (12).

Because a marker with all of the ideal MST characteristics (39) has proved to be elusive, “toolbox” approaches using multiple methods can be useful, recognizing that the strengths of one marker can complement the weaknesses of another marker (1, 40). In this study, we developed a human mtDNA ddPCR assay, hCYTB484, and compared its performance characteristics with a widely used microbial DNA marker, HF183/BacR287. The hCYTB484 marker was highly specific to human feces and quantifiable in each of the tested human feces from the US, Bangladesh, and Mozambique, whereas the HF183/BacR287 marker was less widely distributed. In light of the recommendations for validations of microbial MST markers across various geographies, aspects of mtDNA MST methods such as the direct targeting of host DNA and the wide distribution of mtDNA markers throughout individuals illuminate mtDNA methods as a useful set of tools in the MST toolbox. However, other characteristics of mtDNA MST markers such as overall concentration across a range of relevant waste streams, relative persistence in environmental media (41, 42), cellular origins of fecal mtDNA, relationship with pathogen fate and transport in environmental media, and relationship with risk have not been well characterized and warrant further study due to their relevance in environmental monitoring. As an unambiguous indicator of human source, mtDNA may complement other markers that have been better characterized in terms of the aforementioned unknowns but have been limited in source-specificity (10).

### Limitations

Several limitations qualify the results of this study. Our specificity testing was limited in the diversity of non-human species of feces tested. While we only used cow and pig feces for testing specificity, we sought to include large sample sets of two of the most relevant non-human sources of fecal pollution in practice. While our aLoD experiments show agreement between theoretical calculations and observed ddPCR experiments, our choice of using the Poisson distribution to calculate our aLoD represents a theoretical aLoD that does not account for biases induced by the operator or equipment. Interpolation approaches to the determination of aLLoQ on are helpful in that they may provide a more standardized approach; however, because the imprecision profiles of ddPCR platforms has not been widely studied, the selection of a model for imprecision profile is not well guided by the literature. Our choice of a linear regression on log_10_ transformed values of both CV and the copies per ddPCR well was guided by visual assessment of our data and the desire to use a regression method widely familiar to others.

## Acknowledgements

This material is based upon work supported by the National Science Foundation under Grant No. 1511825. The funders had no role in study design, data collection and interpretation, or the decision to submit the work for publication. We thank Christine Moe and Amy Pickering for their donation of the US and Bangladesh fecal samples, and we acknowledge Janet Hatt, Trent Sumner, Sid Patel, Victoria Dean, Alexandra Bogožalec Košir, and Stefan Grebe.

## Materials and Methods

### Assay design aims

We aimed to design an assay targeting human mtDNA with three main goals: 1) specificity to human mtDNA, 2) optimization for ddPCR conditions, and 3) avoidance of identified polymorphisms.

### Assay target gene

We chose to design our assay within the cytochrome *b* region on the mtDNA because this region has been used to differentiate between species in other fields of study such as species conservation (43), identification of protected species (44), and meat identification (45). Crucially, the cytochrome *b* gene has been shown to provide species-specific sequences even in short fragments, 358 bp and 148 bp (18, 19, respectively), making this gene an appealing target for relatively short sequence length applications such as qPCR (45).

### Designing for human specificity

We downloaded complete cytochrome *b* sequences from GenBank (48) (see Supplemental Materials for accession numbers) and produced a consensus sequence for human, pig, cow, goat, chicken, dog, rat, and white-tailed deer using Jalview (2.10.3). From the consensus sequences, we performed an alignment of the consensus sequences using Mafft (using the L-INS-i algorithm). We selected a region with high differentiation between species guided by the species-variable regions identified in (47), and then entered that region into the Integrated DNA Technologies, Inc. (IDT, Coralville, Iowa) PrimerQuest online tool along with ddPCR conditions specified by Bio-Rad to design forward primer, reverse primer, and probe sequences.

Once PrimerQuest output a set of designed assays, we searched the IDT-designed primer and probe sequences against human and non-human mtDNA sequences downloaded from GenBank to evaluate the *in-silico* specificity of the sequences using the NCBI BLAST tool (49). When searching against all human mtDNA sequences downloaded from NCBI at the time (March 2018), we looked for oligonucleotide sequences that had the most returns with 100% identity and full coverage across the oligonucleotide sequence. When searching against downloaded non-human mtDNA sequences, we looked for primer sequences that had the fewest returns with the lowest percent identity and emphasized a lack of identity at the 3’ end of the primers to maximize specificity to humans (50).

### Designing for ddPCR conditions

Unlike qPCR, where the signal of individual oligonucleotide-template complex inefficiencies can be overshadowed by the signal from the bulk reaction, ddPCR measures a signal from each individual partition, meaning that variations in PCR efficiencies can be resolved at or near (depending on the copies per droplet number) the single copy level (51). Consequently, we used the following settings as detailed by the Bio-Rad QX200 ddPCR Application Guide when designing the assay: primer concentration at 900 nM, probe concentration at 250 nM, concentration of divalent cations at 3.8 mM, and the concentration of dNTPs at 0.8 mM.

### Avoiding polymorphisms

We identified two frequently described polymorphisms within the region we were designing in: the 15301 and 15326 positions (20–24), relative to the revised Cambridge Reference Sequence, rCRS (52) and avoided these two polymorphisms during the primer and probe design process by proceeding with primers and probe sets that did not include these two positions.

### Feces samples

To evaluate the sensitivity of the assay, we used both human fecal samples and human wastewater samples. We obtained feces samples from the US, Bangladesh, and Mozambique. From the US, we obtained healthy, pre-challenge feces samples from 11 adults (18 to 50 years of age) enrolled in a norovirus challenge study (trial registration number: NCT00674336). From Mozambique, we obtained pre-intervention feces samples from 71 children under 4 years of age enrolled in an urban sanitation intervention study (trial registration number: NCT02362932). From Bangladesh, we obtained feces samples from 142 children under 5 years of age enrolled in a water treatment intervention study (trial registration number: NCT02606981). We stored all feces samples at −80°C prior to nucleic acid extraction. To evaluate the specificity of the assay, we collected 22 cow and 35 pig fresh fecal samples from farms located in north Georgia using sterile 15 mL tubes; we transported and stored samples in a 1:1 mixture with Cary-Blair media.

### Wastewater samples

We obtained a variety of wastewater samples from the Atlanta metropolitan region, including 3 primary influent samples and 1 primary effluent sample from plants ranging in capacity from 16 MGD to 90 MGD. We collected approximately 1 L of each wastewater sample and transported wastewater samples on ice, filtering the samples within 12 hours of collection. Using vacuum filtration, we filtered each sample in duplicate, with volumes ranging from 25 mL to 50 mL of sample onto 0.22 μm pore size, 47 mm diameter, asymmetrical polyethersulfone membrane filters. We then stored the filters at −80°C until nucleic acid extraction.

### Nucleic acids extractions

We extracted the human feces samples from the US using the MoBio PowerSoil kit according to the manufacturer’s protocol and stored eluted DNA in Buffer C6 at −20°C until analysis. We extracted feces samples from Mozambique and Bangladesh using the Qiagen QIAamp 96 PowerFecal QIAcube HT kit on the QIAcube HT platform and stored eluted DNA in Buffer ATE at −80°C in low-retention microcentrifuge tubes until analysis. For all human feces samples, we started extractions by gently mixing the feces sample with a sterile inoculating loop and extracting from approximately 0.1 grams of feces sample. We extracted cow and pig feces samples using the Qiagen PowerSoil kit according to the Human Gut Microbiome Project protocol and stored eluted DNA in Buffer C6 at −80°C in low-retention microcentrifuge tubes until analysis. We assayed each feces extract with the Qubit™ dsDNA HS assay (Invitrogen™) to quantify dsDNA concentration. We extracted DNA from the wastewater sample filters using the Qiagen PowerWater kit according to the manufacturer’s protocol. We performed extraction blanks with each set of extraction and tested each extraction blank with the Qubit dsDNA HS assay, hCYTB484 assay, and HF183/BacR287 assay.

To assess sampling variability within the human feces samples, we extracted a subset (7%) of the human feces samples in duplicate. For the hCYTB484 assay, we calculated a mean percent difference between biological duplicates of 24% (standard deviation of 27%) and observed no changes in detection status (all biological replicates were quantifiable). For the HF183/BacR287 assay, we observed two cases of changes in detection status between biological replicates.

### End-Point PCR amplifications

We used end-point PCR to confirm the presence of host mtDNA for the cow and pig feces samples before testing for specificity—only cow and pig samples with confirmed host mtDNA were tested in this study. We also used end-point PCR to generate PCR product for cloning prior to sequencing in the verification of positive standards and for confirmation of target sequences. Lastly, for each assay, we checked for the presence of primer dimers for each assay by visualizing end-point PCR products on a fluorescent gel; we did not detect presence of primer dimers for both assays. For each 20 μL end-point PCR reaction, we used 10 μL TaKaRa *Premix Taq* 2x concentration, with 300 nM of each primer, and 2 μL of template. We used a thermocycling routine consisting of 98°C for 10 seconds, 55°C for 30 seconds, 72°C for 60 seconds, and 72°C for 7 minutes. For visualization, we used 1% GelRed stain from Biotium (Fremont, CA) with a 2.5% agarose gel and a 20 bp molecular ruler from Bio-Rad Laboratories, Inc. (Hercules, California).

### ddPCR experiments

In reporting the methods and materials for this study, we followed the Minimum Information for Publication of Quantitative Digital PCR Experiments Guidelines (53). We performed our ddPCR experiments on a Bio-Rad QX200™ Droplet Digital™ PCR System in our laboratory. For each ddPCR well, we created a 21 μL PCR mixture containing 1x Bio-Rad ddPCR™ Supermix for Probes, primer (1000 nM for HF183/BacR287 primers and 900 nM for hCYTB484 primers), 250 nM of probe (for HF183/BacR287 probe, HF183/BacR87 IAC probe, and hCYTB484 probe), and 2 μL of template. We did not use additives in the reaction mixture or enzymatic digestion on samples. We performed droplet generation using 20 μL of reaction mixture and 70 μL of Bio-Rad Droplet Generation Oil for Probes with the Bio-Rad QX200 Droplet Generator. We then transferred the oil emulsion using a multichannel pipettor to a ddPCR™ 96-Well Plate. After sealing the plate using the Bio-Rad PX1 PCR Plate Sealer with a foil PCR Plate Heat Seal, we thermocycled each plate using the Bio-Rad C1000 Touch™ Thermal Cycler. To read each plate, we used the Bio-Rad QX200 Droplet Reader set to the absolute quantification experiment setting. We did not measure our partition volumes (54), but utilized the assumed partition volume of 0.85 nL in Bio-Rad QuantaSoft (Version 1.7.4.0917) for concentration calculations.

We ordered all primers (standard desalting purification) used in this study as custom-manufactured DNA oligos from IDT (Coralville, Iowa). We obtained the hCYTB484 probe as a custom-manufactured probe (HPLC purification) with Internal ZEN™ Quencher Placement from IDT. We ordered both probes (BacP234MGB and BacP234IAC) used for the HF183/BacR287 assay from Applied Biosystems™ (Waltham, Massachusetts) as TaqMan® minor groove binder (MGB) probes (HPLC purification).

For each assay, we conducted a series of experiments to optimize the annealing temperature. First, we ran a temperature gradient spanning approximately 8°C; then, we ran a finer scale temperature gradient (spanning approximately 2°C) by identifying the highest temperatures in which the separation between negative and positive bands reached a limit in the previous gradient. We selected an annealing temperature from the finer scale gradient that gave us the most separation while remaining a relatively high temperature to avoid non-specific amplification. We also experimented with 94°C, 95°C, and 96°C denaturation cycles, finding that 95°C worked the best for the assays used in this study. For each thermocycling routine, we used 10 minutes at 95°C, followed by 40 cycles of 30 seconds at 95°C and 60 seconds at the assay-specific annealing temperature, followed by a 10-minute hold at 98°C. We used an annealing temperature of 58°C for HF183/BacR287 and 59°C for hCYTB484. We set the ramp rate of the thermocycling routines at 2°C/second.

We utilized a positive control containing a sequence-verified, positive control for each assay target on each ddPCR plate as explained below. We used two different kinds of positive controls in our analytical limits of detection. For the HF183/BacR287 assay, we constructed plasmids from positive human feces samples, cloned using an Invitrogen™ (Carlsbad, California) TOPO^®^TA Cloning^®^ kit with the pCR™4-TOPO vector, and verified the plasmid sequences using Sanger sequencing (GENEWIZ, South Plainfield, New Jersey). Prior to use in ddPCR, we digested plasmids using EcoRI (Invitrogen™ Anza Restriction Enzyme). For the hCYTB484 assay, we ordered custom-manufactured, sequence-verified synthesized DNA (gBlock) products manufactured by IDT (Coralville, Iowa) consisting of the hCYTB484 amplicon plus flanking randomized padding. The sequence of the positive controls is shown in Table 4. We used the internal amplification control (IAC) as described in Green et al. 2014 to assess PCR inhibition in each sample. We ordered a custom-manufactured, sequence-verified synthesized DNA (gBlock) from IDT for the IAC. We utilized the IAC probe in multiplex each time the HF183/BacR287 assay was run. We also monitored the no-template controls (UV-treated molecular grade water) run with each plate for amplification. If amplification did occur, we re-ran the samples until we obtained a plate with zero positive partitions in the no-template controls. We used undiluted samples for with HF183/BacR287 to monitor each sample for PCR inhibitors using the IAC as described (2). If any sample exceeded the ddPCR dynamic range, we reran it with increasing ten-fold dilutions until the run had a ddPCR λ (mean copies per partition) under 6 and was above the aLLoQ.

**Table 4.**
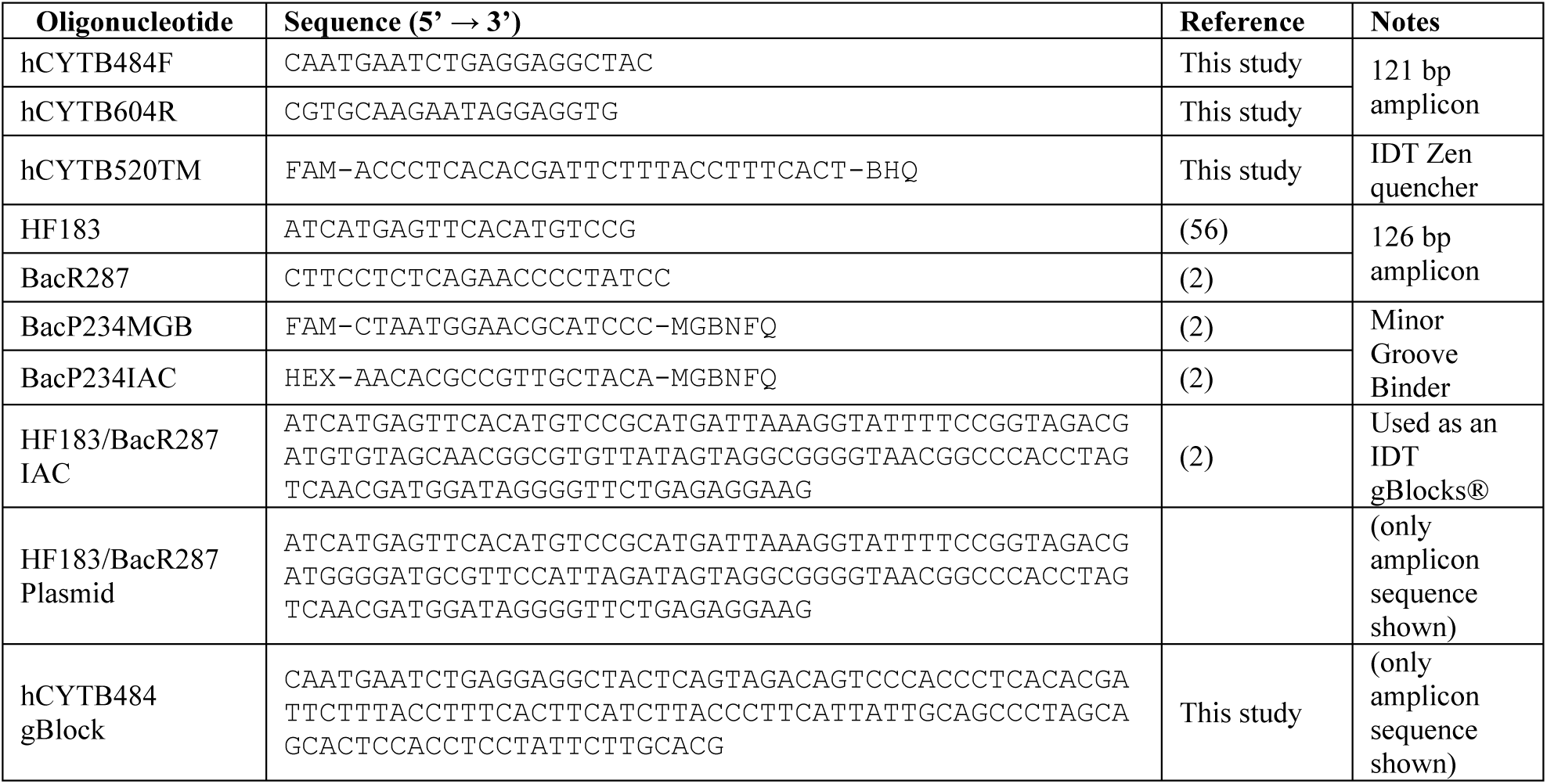
Primers, probes, and positive controls utilized in this study.

Due to previous experience with ddPCR rain, specifically, in situations where primer and template were mismatched by one or two base pairs, we chose to adopt a moderately conservative approach to distinguishing between positive and negative droplets by setting the threshold midway between the negative droplet peak and the positive droplet peak. We intended for this midway setting to avoid including instances of primer and template binding of very similar but not exact sequences as positive partitions. We only accepted ddPCR results from wells with more than 10,000 accepted droplets (histogram of accepted droplets in Figure S4). Our mean lambda (copies per droplet) value was 2.19. We normalized marker counts in feces to ng of dsDNA to account for variability between feces consistencies (55).

### Analytical limits

In order to set the limit of detection and limit of quantification, we selected specific definitions for each limit and performed corresponding ddPCR experiments to validate each definition. A core assumption of our approach is that the number of positive partitions per well can be modeled using the Poisson distribution (27).

### Fitting a Poisson distribution

To model a theoretical Poisson distribution of positive partitions in a ddPCR well corresponding to observed data from ddPCR experiments, we used maximum likelihood estimation (MLE). Assuming that the conditions under which each ddPCR well was prepared, thermocycled, and quantified were equal such that all wells from a particular experiment can be considered to be from the same population, we modeled a Poisson distribution from our observed data. Because the Poisson distribution originates from the exponential family, it can be shown that the MLE is the sample mean (see supplemental materials “Poisson Maximum Likelihood Estimation”).

### Analytical limit of detection

When testing analytical blanks, we assayed UV treated (for 15 minutes) molecular grade water in 94 replicates with both assays (HF183/BacR287 and hCYTB484). We chose to define the limit of detection as the minimum level of the analyte in a sample that will be reported as detected with 95% probability. In order to assess the minimum level of analyte to be detected 95% of the time, we used the Poisson distribution to model the number of positive partitions in a ddPCR well. To determine the level of analyte that would be detected 95% of the time in a Poisson process, we set the Poisson probability mass function equal to a probability of 0.05 for observing no positive partitions in a well.

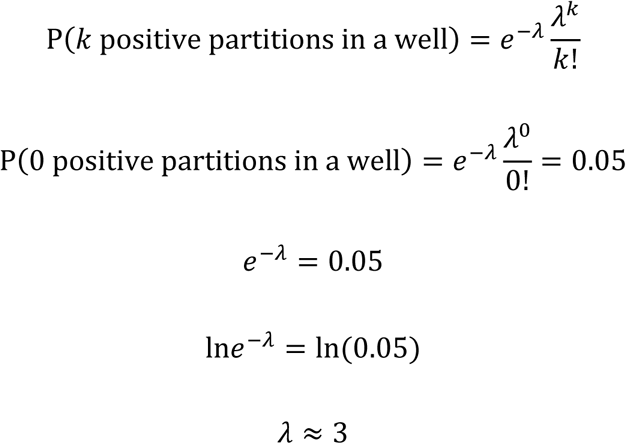

For both assays, we used serial dilutions to dilute the respective positive controls down to approximately 3 positive partitions per well and then assayed the dilution closest to 3 positive partitions per well for 46 ddPCR replicates. This aLoD of 3 positive partitions per well has been previously utilized with applications of ddPCR in the MST field (16) and is recognized as a theoretical aLoD for molecular analyses, such as qPCR (28).

### Analytical lower limit of quantification

We chose to define the analytical lower limit of quantification as the minimum level of analyte in a sample that will result in a coefficient of variation (CV) of 25%. The target CV of 25% or less is a commonly used aLLoQ goal for ddPCR platforms (30, 31). The CV, also referred to as the relative standard deviation (RSD), is defined as the ratio of the standard deviation, σ, to the mean, μ.

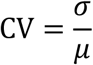

Because repeated assaying of various target concentrations may be required to meet a specific CV target, we sought to understand how the assay imprecision changes with level of concentration to allow us to interpolate estimates of the level of concentrations needed to meet our target CV. In an approach similar to that used in Milosevic et al. 2018 (31), we assayed a variety of concentration levels, calculated CV values, and fitted linear regressions to observed CV values plotted against the concentration of target in a ddPCR well (copies per 20 μL well). However, to be able to interpolate the level of concentration needed to reach our target CV using the regression, we needed to estimate the true CV value so that regressions resulting from calculated CV values are truly reflective of the assay’s imprecision profile and not an artifact of too few replicates (29). Thus, we used a relatively large number of replicates in conjunction with cumulative moving plots of the CV. The cumulative moving plots of the CV served to allow us to see if the cumulative moving CV converged before the total number of replicates. Ideally, if we had more than enough replicates for the estimation of the true CV, the cumulative moving CV plot would converge on the CV value calculated from the total number of replicates and remain there (Figure S3H). Once we estimated CV values for a variety of concentrations, we log-transformed both the CV values and concentrations of target in a ddPCR well and fit a linear regression.

### Empirical estimation of coefficients of variation

Because the number of ddPCR replicates used to estimate the CV of a levels of concentration varies in the literature (30, 31), we used cumulative moving plots of the CV for each sample set, plotting the CV as a function of the number of replicates. We used these cumulative moving plots to visualize when the CV as a function of number of replicates converged around the mean CV of the total sample set.

### Sensitivity and specificity testing

Prior to assaying the cow and pig feces samples with hCYTB484 and HF183/BacR287, we confirmed each sample for the presence of the respective host-animal mtDNA (13) using end-point PCR. After confirmation, we ran both assays on each cow and pig feces sample in duplicate on ddPCR. We Sanger sequenced (GENEWIZ, South Plainfield, NJ) each sample that was above the aLoD the end-point PCR product. We confirmed that each of the 3 sequenced HF183/BacR287 positive samples contained the HF183/BacR287 standard sequence. We did not confirm the presence of the hCYTB484 target sequence in the 3 hCYTB484 positive samples, nor the presence of the respective animal host mtDNA. We performed further testing with other human-specific mtDNA assays (13, 14) and observed amplification below the aLoD. This suggests the potential for human mtDNA contamination of the cow and pig samples as the source of amplification; however, we chose to regard these samples as false positive for hCYTB484 to be conservative. We assayed each human feces sample using hCYTB484 and HF183/BacR287 on ddPCR with 25% technical replicates.

### Statistical analyses

All statistical analyses, simulations, and graphs were done in R language, using R version 3.6.1.

## Supplemental Material

### Figures

**Figure S1.**
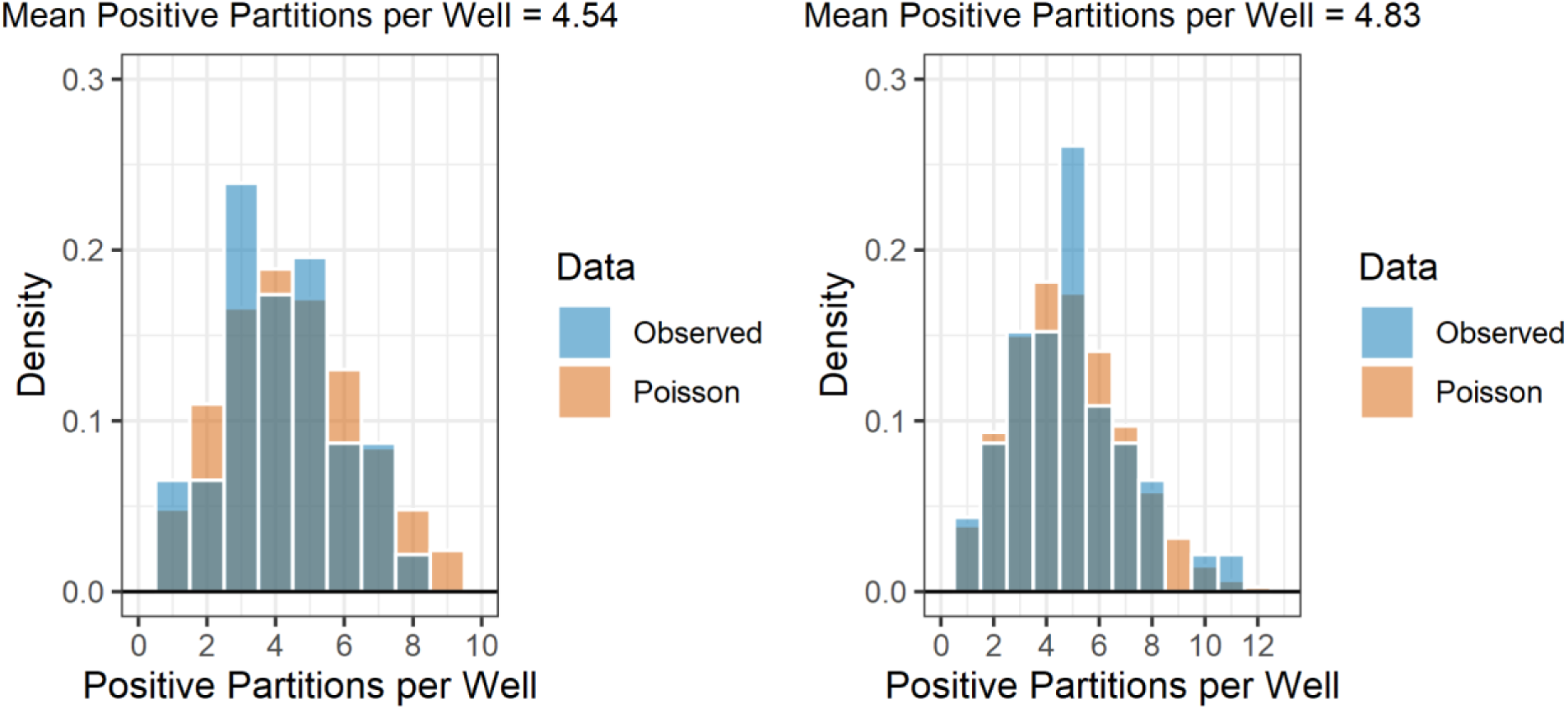
Comparisons of sampling distributions with corresponding expected Poisson distributions generated using maximum likelihood estimation. Left and right plots show results from the hCYTB484 and HF183/BacR287 assays, respectively. *P*-values from the results of Anderson-Darling tests between theoretical and observed distributions are shown in the top right corners of each plot.

**Figure S2.**
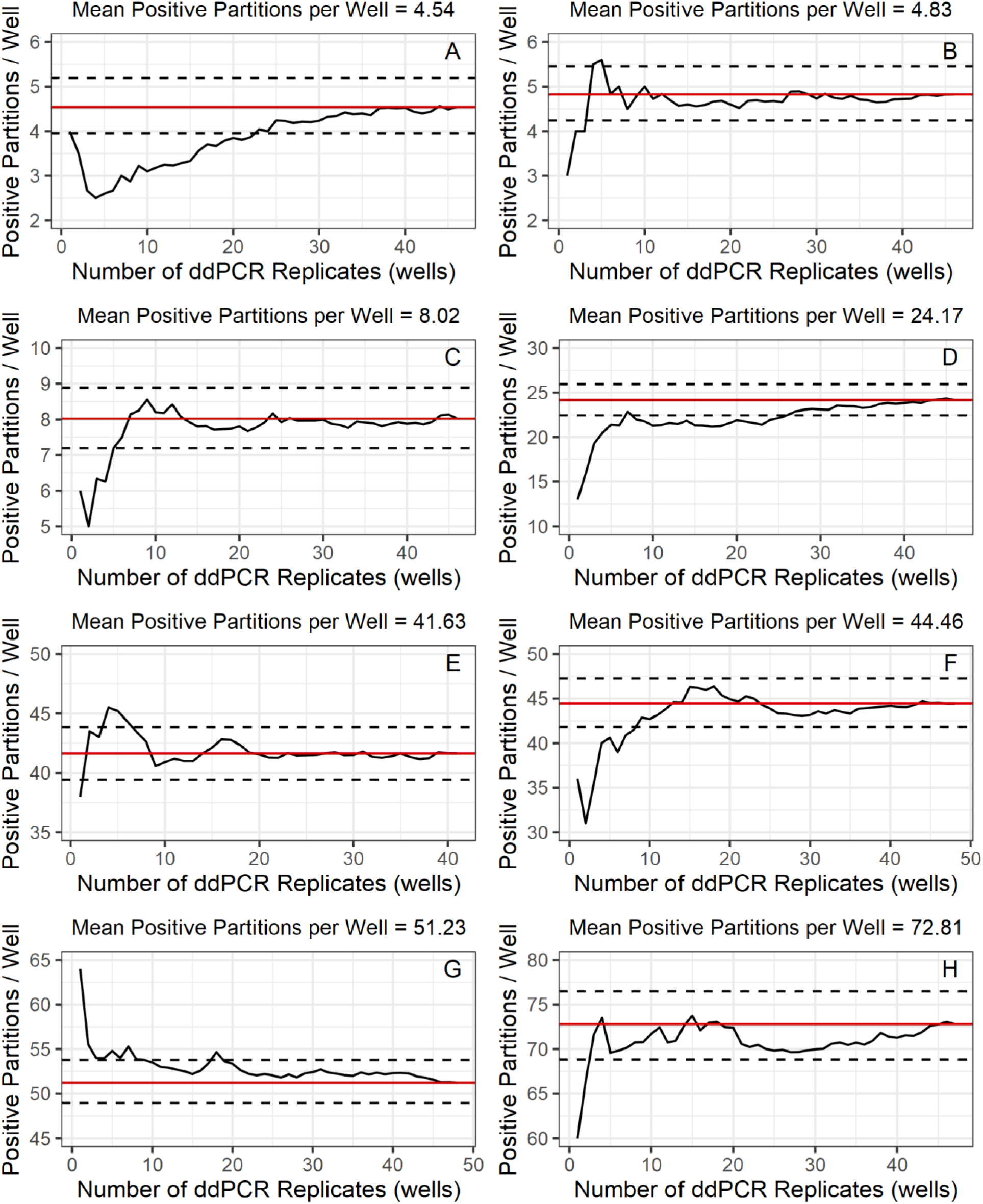
Cumulative moving plots of mean positive partitions per well for a range of mean positive partitions per well. The mean positive partitions per well calculated from total number of replicates is shown as the red solid horizontal line and bootstrapped 95% confidence interval is shown as the black dotted horizontal lines.

**Figure S3.**
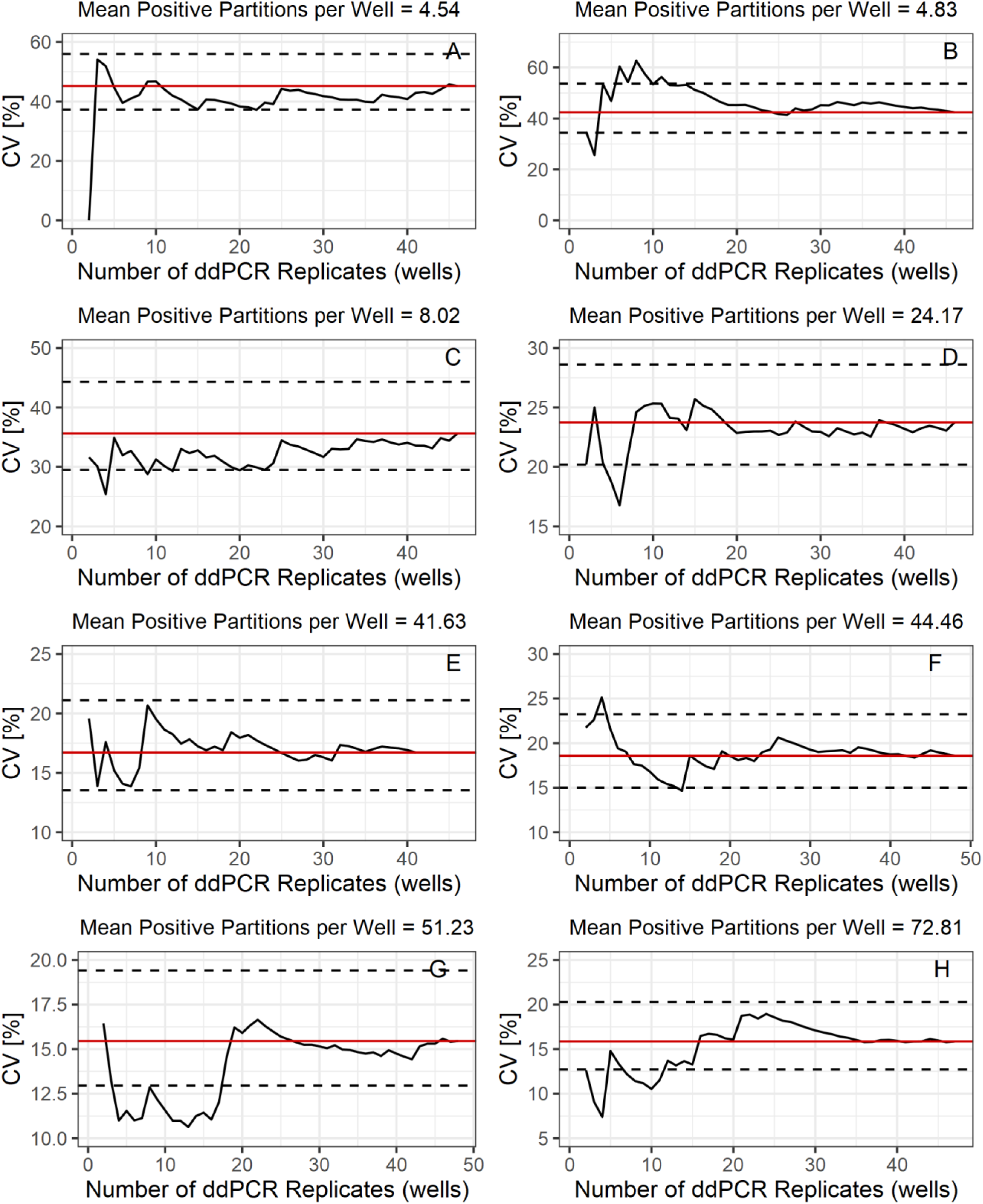
Cumulative moving plots of CV for a range of mean positive partitions per well. The CV calculated from total number of replicates is shown as the red solid horizontal line and bootstrapped 95% confidence interval is shown as the black dotted horizontal lines.

**Figure S4.**
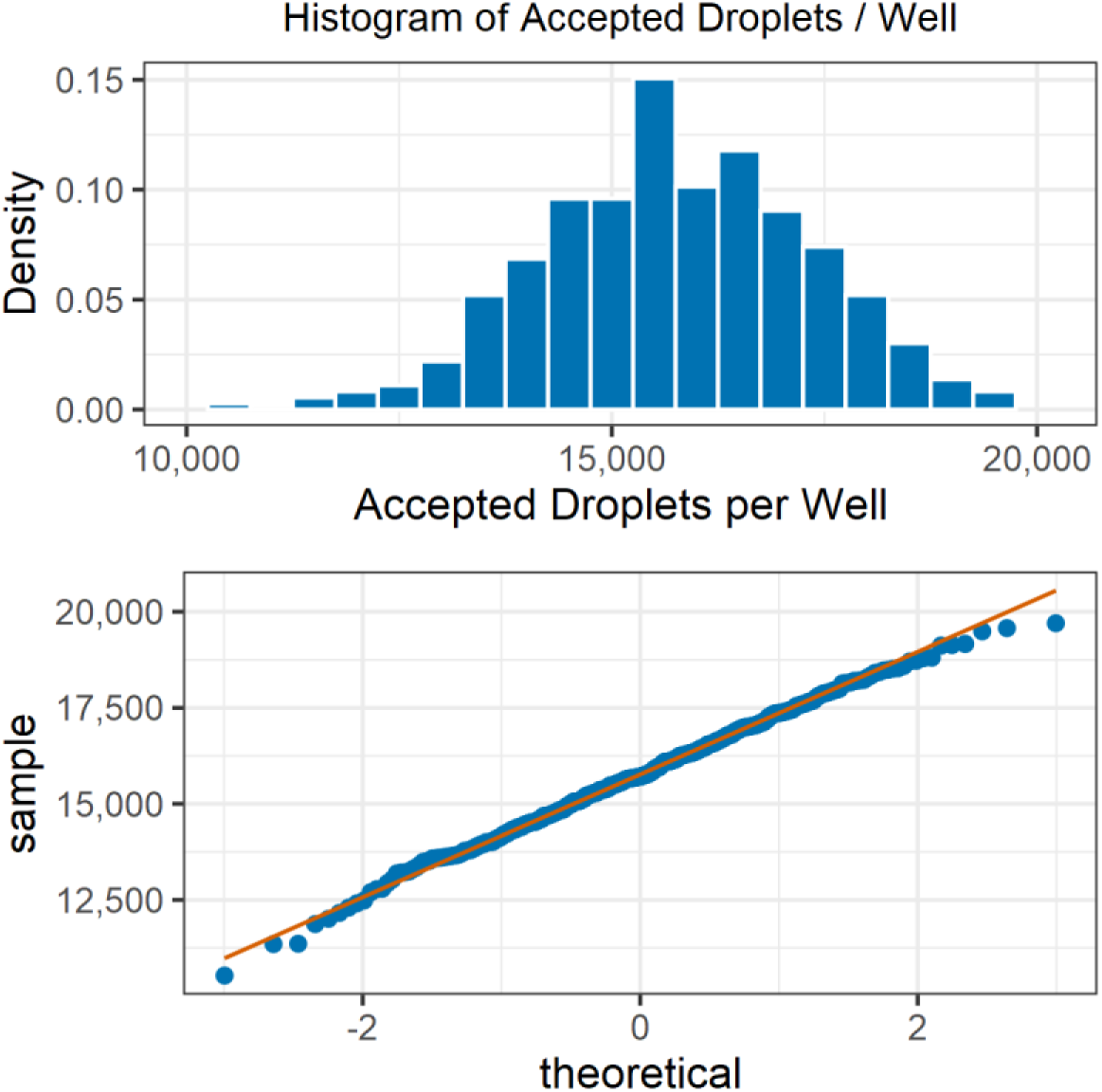
Histogram of the accepted droplets (top) and quantile-quantile plot comparing the distribution of accepted droplets with a theoretical normal distribution along with a quantile-quantile line for comparison (bottom). These plots show the numbers of accepted droplets for the ddPCR experiments conducted in this study on the Bio-Rad QX200 platform.

## Poisson Maximum Likelihood Estimation

Suppose X_1_,…,X_n_ are random samples from a Poisson distribution with parameter **λ**. To derive the maximum likelihood estimator, we start with the Poisson probability mass function:

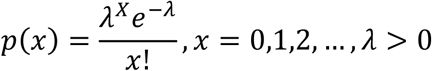

Likelihood function:

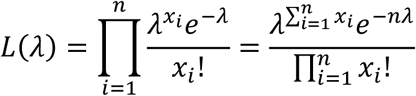

Taking the natural logarithm:

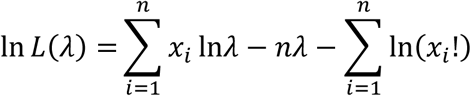

Differentiating with respect to λ:

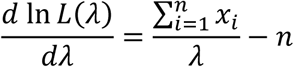

Setting the derivative (right hand side) equal to zero:

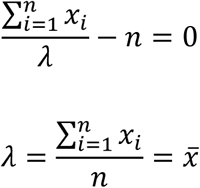

Therefore, the MLE of λ is

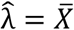

## References

1. Boehm AB, Van De Werfhorst LC, Griffith JF, Holden PA, Jay JA, Shanks OC, Wang D, Weisberg SB. 2013. Performance of forty-one microbial source tracking methods: A twenty-seven lab evaluation study. Water Res 47:6812–6828.

2. Green HC, Haugland RA, Varma M, Millen HT, Borchardt MA, Field KG, Walters WA, Knight R, Sivaganesan M, Kelty CA, Shanks OC. 2014. Improved HF183 Quantitative Real-Time PCR Assay for Characterization of Human Fecal Pollution in Ambient Surface Water Samples. Appl Environ Microbiol 80:3086–3094.

3. Stachler E, Kelty C, Sivaganesan M, Li X, Bibby K, Shanks OC. 2017. Quantitative CrAssphage PCR Assays for Human Fecal Pollution Measurement. Environ Sci Technol 51:9146–9154.

4. US EPA. 2019. Method 1697: Characterization of Human Fecal Pollution in Water by Polymerase Chain Reaction (qPCR) Assay.

5. Harris AR, Pickering AJ, Harris M, Doza S, Islam MS, Unicomb L, Luby S, Davis J, Boehm AB. 2016. Ruminants Contribute Fecal Contamination to the Urban Household Environment in Dhaka, Bangladesh. Environ Sci Technol 50:4642–4649.

6. Nshimyimana JP, Cruz MC, Thompson RJ, Wuertz S. 2017. Bacteroidales markers for microbial source tracking in Southeast Asia. Water Res 118:239–248.

7. Reischer GH, Ebdon JE, Bauer JM, Schuster N, Ahmed W, Åström J, Blanch AR, Blöschl G, Byamukama D, Coakley T, Ferguson C, Goshu G, Ko G, De Roda Husman AM, Mushi D, Poma R, Pradhan B, Rajal V, Schade MA, Sommer R, Taylor H, Toth EM, Vrajmasu V, Wuertz S, MacH RL, Farnleitner AH. 2013. Performance characteristics of qPCR assays targeting human- and ruminant-associated bacteroidetes for microbial source tracking across sixteen countries on six continents. Environ Sci Technol 47:8548–8556.

8. Odagiri M, Schriewer A, Hanley K, Wuertz S, Misra PR, Panigrahi P, Jenkins MW. 2015. Validation of Bacteroidales quantitative PCR assays targeting human and animal fecal contamination in the public and domestic domains in India. Sci Total Environ 502:462– 470.

9. Yatsunenko T, Rey FE, Manary MJ, Trehan I, Dominguez-Bello MG, Contreras M, Magris M, Hidalgo G, Baldassano RN, Anokhin AP, Heath AC, Warner B, Reeder J, Kuczynski J, Caporaso JG, Lozupone CA, Lauber C, Clemente JC, Knights D, Knight R, Gordon JI. 2012. Human gut microbiome viewed across age and geography. Nature 486:222–7.

10. Mayer RE, Reischer GH, Ixenmaier SK, Derx J, Blaschke AP, Ebdon JE, Linke R, Egle L, Ahmed W, Blanch AR, Byamukama D, Savill M, Mushi D, Cristóbal HA, Edge TA, Schade MA, Aslan A, Brooks YM, Sommer R, Masago Y, Sato MI, Taylor HD, Rose JB, Wuertz S, Shanks OC, Piringer H, Mach RL, Savio D, Zessner M, Farnleitner AH. 2018. Global Distribution of Human-Associated Fecal Genetic Markers in Reference Samples from Six Continents. Environ Sci Technol 52:5076–5084.

11. Harris AAR, Pickering AJ, Harris M, Doza S, Islam S, Unicomb L, Luby S, Davis J, Alexandria B. 2016. Ruminants contribute fecal contamination to the urban household environment in Dhaka, Bangladesh 1–15.

12. Berendes DM, Yang PJ, Lai A, Hu D, Brown J. 2018. Estimation of global recoverable human and animal faecal biomass. Nat Sustain 1:679–685.

13. Caldwell JM, Raley ME, Levine JF. 2007. Mitochondrial Multiplex Real-Time PCR as a Source Tracking Method in Fecal-Contaminated Effluents. Environ Sci Technol 41:3277– 3283.

14. Schill WB, Mathes M V. 2008. Real-Time PCR Detection and Quantification of Nine Potential Sources of Fecal Contamination by Analysis of Mitochondrial Cytochrome b Targets. Environ Sci Technol 42:5229–5234.

15. He X, Liu P, Zheng G, Chen H, Shi W, Cui Y, Ren H. 2016. Evaluation of five microbial and four mitochondrial DNA markers for tracking human and pig fecal pollution in freshwater. Nat Publ Gr 1–11.

16. Cao Y, Raith MR, Griffith JF. 2015. Droplet digital PCR for simultaneous quantification of general and human-associated fecal indicators for water quality assessment. Water Res 70:337–349.

17. Hindson CM, Chevillet JR, Briggs HA, Gallichotte EN, Ruf IK, Hindson BJ, Vessella RL, Tewari M. 2013. Absolute quantification by droplet digital PCR versus analog real-time PCR. Nat Methods 10:1003–1005.

18. Strain MC, Lada SM, Luong T, Rought SE, Gianella S, Terry VH, Spina CA, Woelk CH, Richman DD. 2013. Highly Precise Measurement of HIV DNA by Droplet Digital PCR. PLoS One 8:2–9.

19. Stadhouders R, Pas SD, Anber J, Voermans J, Mes THM, Schutten M. 2010. The Effect of Primer-Template Mismatches on the Detection and Quantification of Nucleic Acids Using the 5′ Nuclease Assay. J Mol Diagnostics 12:109–117.

20. Ablimit A, Qin W, Shan W, Wu W, Ling F, Ling KH, Zhao C, Zhang F, Ma Z, Zheng X. 2013. Genetic diversities of cytochrome B in Xinjiang Uyghur unveiled its origin and migration history. BMC Genet 14:1.

21. Hwa HL, Ko TM, Chen YC, Chang YY, Tseng LH, Su YN, Lee JCI. 2010. Study of the cytochrome b gene sequence in populations of Taiwan. J Forensic Sci 55:167–170.

22. Lee SD, Lee YS, Lee J Bin. 2002. Polymorphism in the mitochondrial cytochrome B gene in Koreans. Int J Legal Med 116:74–78.

23. Amer SAM, Alhothali B, Tubaigy SMA. 2015. Possible application of cytochrome b gene for human identification. Arab J Forensic Sci Forensic Med 1:24–248.

24. Farghadani R, Babadi AA. 2015. Nucleotide variation of the mitochondrial cytochrome b gene in the Malay population. Rom J Leg Med 23:57–60.

25. Letowski J, Brousseau R, Masson L. 2004. Designing better probes: effect of probe size, mismatch position and number on hybridization in DNA oligonucleotide microarrays. J Microbiol Methods 57:269–278.

26. Haugland RA, Varma M, Sivaganesan M, Kelty C, Peed L, Shanks OC. 2010. Evaluation of genetic markers from the 16S rRNA gene V2 region for use in quantitative detection of selected Bacteroidales species and human fecal waste by qPCR. Syst Appl Microbiol 33:348–357.

27. 2018. Digital PCR. Springer New York, New York, NY.

28. Bustin SA, Benes V, Garson JA, Hellemans J, Huggett J, Kubista M, Mueller R, Nolan T, Pfaffl MW, Shipley GL, Vandesompele J, Wittwer CT. 2009. The MIQE guidelines: Minimum information for publication of quantitative real-time PCR experiments. Clin Chem 55:611–622.

29. Forootan A, Sjöback R, Björkman J, Sjögreen B, Linz L, Kubista M. 2017. Methods to determine limit of detection and limit of quantification in quantitative real-time PCR (qPCR). Biomol Detect Quantif 12:1–6.

30. Bogožalec Košir A, Demšar T, Štebih D, Žel J, Milavec M. 2019. Digital PCR as an effective tool for GMO quantification in complex matrices. Food Chem 294:73–78.

31. Milosevic D, Mills JR, Campion MB, Vidal-Folch N, Voss JS, Halling KC, Highsmith WE, Liu MC, Kipp BR, Grebe SKG. 2018. Applying standard clinical chemistry assay validation to droplet digital PCR quantitative liquid biopsy testing. Clin Chem 64:1732– 1742.

32. Dobnik D, Spilsberg B, Bogožalec Košir A, Holst-Jensen A, Žel J. 2015. Multiplex Quantification of 12 European Union Authorized Genetically Modified Maize Lines with Droplet Digital Polymerase Chain Reaction. Anal Chem 87:8218–8226.

33. Sadler WA. 2016. Using the variance function to estimate limit of blank, limit of detection and their confidence intervals. Ann Clin Biochem 53:141–149.

34. Davidian M, Carroll RJ. 1987. Variance function estimation. J Am Stat Assoc 82:1079– 1091.

35. Berweger CD, Müller-Plathe F, Hänseler E, Keller H. 1998. Estimating imprecision profiles in biochemical analysis. Clin Chim Acta 277:107–125.

36. Fawcett T. 2006. An introduction to ROC analysis. Pattern Recognit Lett 27:861–874.

37. Morrison AM, Coughlin K, Shine JP, Coull BA, Rex AC. 2003. Receiver Operating Characteristic Curve Analysis of Beach Water Quality Indicator Variables. Appl Environ Microbiol 69:6405–6411.

38. Layton BA, Cao Y, Ebentier DL, Hanley K, Ballesté E, Brandão J, Byappanahalli M, Converse R, Farnleitner AH, Gentry-Shields J, Gidley ML, Gourmelon M, Lee CS, Lee J, Lozach S, Madi T, Meijer WG, Noble R, Peed L, Reischer GH, Rodrigues R, Rose JB, Schriewer A, Sinigalliano C, Srinivasan S, Stewart J, Van De Werfhorst LC, Wang D, Whitman R, Wuertz S, Jay J, Holden PA, Boehm AB, Shanks O, Griffith JF. 2013. Performance of human fecal anaerobe-associated PCR-based assays in a multi-laboratory method evaluation study. Water Res 47:6897–6908.

39. 2011. Microbial Source Tracking: Methods, Applications, and Case Studies. Springer New York, New York, NY.

40. Harwood VJ, Staley C, Badgley BD, Borges K, Korajkic A. 2014. Microbial source tracking markers for detection of fecal contamination in environmental waters: Relationships between pathogens and human health outcomes. FEMS Microbiol Rev 38:1–40.

41. He X, Chen H, Shi W, Cui Y, Zhang XX. 2015. Persistence of mitochondrial DNA markers as fecal indicators in water environments. Sci Total Environ 533:383–390.

42. Zimmerman BD, Ashbolt NJ, Garland JL, Keely S, Wendell D. 2014. Human mitochondrial DNA and endogenous bacterial surrogates for risk assessment of graywater reuse. Environ Sci Technol 48:7993–8002.

43. Hsieh H-M, Chiang H-L, Tsai L-C, Lai S-Y, Huang N-E, Linacre A, Lee JC-I. 2001. Cytochrome b gene for species identification of the conservation animals. Forensic Sci Int 122:7–18.

44. Lee JCI, Hsieh HM, Huang LH, Kuo YC, Wu JH, Chin SC, Lee AH, Linacre A, Tsai LC. 2009. Ivory identification by DNA profiling of cytochrome b gene. Int J Legal Med 123:117–121.

45. Dooley JJ, Paine KE, Garrett SD, Brown HM. 2004. Detection of meat species using TaqMan real-time PCR assays. Meat Sci 68:431–438.

46. Parson W, Pegoraro K, Niederstätter H, Föger M, Steinlechner M. 2000. Species identification by means of the cytochrome b gene. Int J Legal Med 114:23–28.

47. Lopez-Oceja A, Gamarra D, Borragan S, Jiménez-Moreno S, de Pancorbo MM. 2016. New cyt b gene universal primer set for forensic analysis. Forensic Sci Int Genet 23:159– 165.

48. Benson DA, Cavanaugh M, Clark K, Karsch-Mizrachi I, Lipman DJ, Ostell J, Sayers EW. 2017. GenBank. Nucleic Acids Res 45:D37–D42.

49. Camacho C, Coulouris G, Avagyan V, Ma N, Papadopoulos J, Bealer K, Madden TL. 2009. BLAST+: architecture and applications. BMC Bioinformatics 10:421.

50. Wright ES, Yilmaz LS, Ram S, Gasser JM, Harrington GW, Noguera DR. 2014. Exploiting extension bias in polymerase chain reaction to improve primer specificity in ensembles of nearly identical DNA templates. Environ Microbiol 16:1354–1365.

51. Witte AK, Mester P, Fister S, Witte M, Schoder D, Rossmanith P. 2016. A Systematic Investigation of Parameters Influencing Droplet Rain in the Listeria monocytogenes prfA Assay - Reduction of Ambiguous Results in ddPCR. PLoS One 11:e0168179.

52. Bandelt HJ, Kloss-Brandstätter A, Richards MB, Yao YG, Logan I. 2014. The case for the continuing use of the revised Cambridge Reference Sequence (rCRS) and the standardization of notation in human mitochondrial DNA studies. J Hum Genet 59:66–77.

53. Huggett JF, Foy CA, Benes V, Emslie K, Garson JA, Haynes R, Hellemans J, Kubista M, Mueller RD, Nolan T, Pfaffl MW, Shipley GL, Vandesompele J, Wittwer CT, Bustin SA. 2013. The Digital MIQE Guidelines: Minimum Information for Publication of Quantitative Digital PCR Experiments. Clin Chem 59:892–902.

54. Košir AB, Divieto C, Pavšič J, Pavarelli S, Dobnik D, Dreo T, Bellotti R, Sassi MP, Žel J. 2017. Droplet volume variability as a critical factor for accuracy of absolute quantification using droplet digital PCR. Anal Bioanal Chem 409:6689–6697.

55. Kelty CA, Varma M, Sivaganesan M, Haugland RA, Shanks OC. 2012. Distribution of Genetic Marker Concentrations for Fecal Indicator Bacteria in Sewage and Animal Feces. Appl Environ Microbiol 78:4225–4232.

56. Bernhard AE, Field KG. 2000. A PCR assay to discriminate human and ruminant feces on the basis of host differences in Bacteroides-Prevotella genes encoding 16S rRNA. Appl Environ Microbiol 66:4571–4574.

